# Centrosome Aurora A gradient ensures a single PAR-2 polarity axis by regulating RhoGEF ECT-2 localization in *C. elegans* embryos

**DOI:** 10.1101/396721

**Authors:** Sukriti Kapoor, Sachin Kotak

**Author notes:** MCB (IISc) SE-12 560012 Bangalore India.

## Abstract

The proper establishment of the cell polarity is essential for development and morphogenesis. In the *Caenorhabditis elegans* one-cell embryo, a centrosome localized signal provides spatial information that is responsible for generating a single polarity axis. It is hypothesized that such a signal causes local inhibition of cortical actomyosin network in the vicinity of the centrosome. This pivotal event initiates symmetry breaking to direct partitioning of the partition defective proteins (PARs) in the one-cell embryo. However, the molecular nature of the centrosome regulated signal that impinges on the posterior cortex to bring upon cortical anisotropy in the actomyosin network and to promote polarity establishment remains elusive. Here, we discover that Aurora A kinase (AIR-1 in *C. elegans*) is essential for proper cortical contractility in the one-cell embryo. Loss of AIR-1 causes pronounced cortical contractions on the entire embryo surface during polarity establishment phase, and this creates more than one PAR-2 polarity axis. Moreover, we show that in the absence of AIR-1, centrosome positioning becomes dispensable in dictating the PAR-2 polarity axis. Interestingly, we identify that Rho Guanine Exchange Factor (GEF) ECT-2 acts downstream to AIR-1 to control excess contractility and notably AIR-1 loss affects ECT-2 cortical localization and thereby polarity establishment. Overall, our study unravels a novel insight whereby an evolutionarily conserved kinase Aurora A inhibits promiscuous PAR-2 domain formation and ensures singularity in the polarity establishment axis.

## Introduction

The potential of cells to establish polarity is essential for several events during development and morphogenesis. A myriad of distinct signals are deployed by various cell types to trigger polarity establishment (reviewed in Chant, 1999; Nelson, 2003). The one-cell stage of *Caenorhabditis elegans* embryo has proven to be an excellent system to investigate the mechanisms of polarity establishment. In this remarkable cell, centrioles brought by the fertilization, when gets mature to centrosome play a critical function for symmetry breaking and thus setting up polarity (Cowan and Hyman, 2004; reviewed in Hoege and Hyman, 2013; Motegi and Seydoux, 2013; Rose and Gönczy, 2014). It has been suggested that pericentriolar material (PCM) proteins of the centrosome are needed for polarity establishment (Goldstein and Hird, 1996; O‵Connell et al., 2000; Hamill et al., 2002); furthermore, the microtubules are also linked in setting up proper polarity in the one-cell embryo (Tsai and Ahringer, 2007; reviewed in Siegrist and Doe, 2007).

Functional characterization of several mutants suggest that centrosome cortical distance is correlated with the defects in the polarity establishment (Rappleye et al., 2002; Rappleye et al., 2003; Fortin et al., 2010). Interestingly, a relatively recent study has suggested that centrosomes are capable of initiating polarity from any position within the one-cell zygote and the proximity of the centrosome to the cell cortex is a critical determining factor for defining the positioning and the timing of polarity initiation (Bienkowska and Cowan, 2012). This study also suggests that the centrosome may carry a gradient of a diffusive signal that acts as a molecular ruler to instruct polarity establishment at the closest cell cortex. However, the nature of such a signal that could guide polarity-initiation in the one-cell embryo remains elusive.

How does the centrosome instruct the neighbouring cell cortex to establish cell polarity? It is proposed that the local inhibition of the actomyosin contractions in the vicinity of centrosomes provides the trigger to commence anisotropy in the one-cell embryo (Munro et al., 2004; reviewed in Cowan and Hyman, 2007). This event can lead to the formation of partition defective proteins [aPARs: PAR-3, PAR-6, and atypical protein kinase C (aPKC) and pPARs: PAR-1, PAR-2 and LGL] domains (reviewed in Motegi and Seydoux, 2013; Hoege and Hyman, 2013). Mutants defective for actomyosin-based contractility are impaired in establishing polarity (Guo and Kemphues, 1996; Hill and Strome, 1990; Severson and Bowerman, 2003). It appears that the initiation of contractile asymmetry in the one-cell embryo is independent of the PAR-polarity, for instance, PAR-6 could be asymmetrically distributed at the anterior cortex in the absence of PAR-2 (Cuenca et al., 2003; Goehring et al., 2011). A small GTPase RHO-1 and its activator RhoGEF ECT-2 play a critical role in controlling contractile asymmetry by modulating the actomyosin network (Motegi and Sugimoto, 2006). RNAi-mediated depletion of ECT-2 or its activator NOP-1 cause impairment in cortical contractility (Motegi and Sugimoto, 2006; Tse et al., 2012). In such embryos, the pPAR domain eventually forms possibly because of the microtubule-dependent polarity pathway that kicks in at the time of polarity maintenance (Zonies et al., 2010; Motegi et al., 2011; Tse et al., 2012; reviewed in Motegi and Seydoux, 2013; Rose and Gönczy, 2014). Remarkably, ECT-2 is excluded from the posterior cortex at the onset of symmetry breaking, and this event is correlated with the disappearance of the non-muscle myosin II (NMY-2) from the posterior cortex (Munro et al., 2004; Motegi and Sugimoto, 2006). However, the molecular pathway that impinges on the RhoGEF ECT-2 so that it is excluded from the posterior cortex at the onset of polarity establishment remains unknown.

Depletion of the PP6 phosphatase catalytic subunit PPH-6 or its regulatory subunit SAPS-1 causes a loss of cortical contractility and disappearance of pseudocleavage in the one-cell embryo (Afshar et al., 2010). We have recently shown that SAPS-1 interacts with Aurora A kinase (AIR-1) in the *C. elegans* embryo and the interplay between AIR-1 and SAPS-1 is crucial for mitotic spindle positioning in the one-cell embryo (Kotak et al., 2016). AIR-1 is a serine/threonine kinase that is essential for the timely mitotic entry, centrosome separation, centrosome maturation, spindle assembly, spindle positioning, spindle elongation and cytokinesis (Hannak et al., 2001; Giet et al., 2002; Toji et al., 2004; Tsai and Zheng, 2005; Hachet et al., 2007; Portier et al., 2007 Seki et al., 2008; Wong et al., 2008; Zheng et al., 2008; Reboutier et al., 2013; Kotak et al., 2016; Mangal et al., 2018). Auto-phosphorylation of AIR-1 at Threonine 201 (Threonine 288 in humans) in its activation loop increases the catalytic activity of Aurora A (Walter et al., 2000; Littlepage et al., 2002; Toya et al., 2011). Interestingly, biochemical and cell biological data suggest that in human cells PP6 phosphatase acts as a T-loop phosphatase for T288 of Aurora A and keeps its activity in a balanced state for proper spindle assembly (Zeng et al., 2010).

In the present work, we show that in contrast to PP6 phosphatase, loss of AIR-1 causes excess cortical contractility at the time of polarity initiation and this excessive contractility translates into polarity defects whereby more than one pPAR axis is set up in the one-cell zygote. We reveal that this function of AIR-1 is dependent on its autocatalytic activity but not on its coactivator TPXL-1 (TPX-2 in humans). Our results further suggest that the impact of AIR-1 depletion on proper polarity establishment is independent of its function in regulating microtubule number. Notably, loss of AIR-1 makes the centrosome dispensable in choreographing the positioning of polarity establishment. Moreover, we show that AIR-1 is required to timely exclude RhoGEF ECT-2 from the posterior cortex. Overall, our study reveals the mechanism by which the AIR-1 gradient at the centrosome ensures singularity in the polarity axis by locally excluding ECT-2 at the posterior cortex.

## Results

### Loss of Aurora A kinase (AIR-1) cause excess cortical contractility in the one-cell embryo

PPH-6 phosphatase and its regulatory subunit SAPS-1 influences cortical contractility in the one-cell stage of *C. elegans* embryo during the symmetry breaking phase (referred to as polarity establishment from now onwards) by influencing the cortical organization of the non-muscle myosin II (NMY-2) (compare Figure 1B with 1A; Supplementary Movies S1 and S2; Afshar et al., 2010). We have recently reported that SAPS-1 interacts with Aurora A kinase (AIR-1) in *C. elegans* embryos (Kotak et al., 2016) and therefore, we analyzed the influence of AIR-1 depletion on cortical contractility in the early one-cell embryo at the time of polarity set-up. RNAi-mediated depletion of AIR-1 in the one-cell embryos impacts centrosome maturation, and this results in collapsing of the centrosomes after nuclear envelope breakdown (NEBD) during prophase (Hannak et al., 2001; Supplementary Movie S3). Also, depletion of AIR-1 influences the centrosome localization of TAC-1 (Le Bot et al., 2003), and therefore we measured GFP-TAC-1 centrosome signal as an another means to check for AIR-1 depletion in our experimental setting, and we found that GFP-TAC-1 levels are highly diminished at the centrosomes (Supplementary Figure S1A-S1C). Thus, we chose this experimental regimen as a proxy for the AIR-1 depletion. Notably, RNAi-mediated loss of AIR-1 causes exaggerated cortical contractility at the time of polarity establishment that is the opposite phenotype to that of SAPS-1 depletion (compare Figure 1C with 1A; Supplementary Movies S3). In human cells, PP6 phosphatase de-phosphorylates Aurora A kinase at T288 [T201 in AIR-1; Supplementary Figure S1D; Toya et al., 2011] to modulate Aurora A activity and this is crucial for the proper spindle assembly (Zeng et al., 2010). Therefore, we postulated that SAPS-1 could be modulating AIR-1 activity for proper cortical contractility and if this is the case, we expect that double depletion of AIR-1 and SAPS-1 should mimic the phenotype of AIR-1 loss. Indeed, we observed that depletion of SAPS-1 and AIR-1 by RNAi [depicted as *saps-1 (RNAi); air-1 (RNAi)*] embryos revealed a phenotype similar to *air-1 (RNAi)* (Figure 1D; Supplementary Movie S4), suggesting that AIR-1 acts downstream of PPH-6 phosphatase complex in regulating cortical contractility. These data further indicate that AIR-1 kinase activity could be crucial for maintaining proper cortical contractility. Thus, we sought to examine AIR-1 kinase function for this phenotype. To this end, we monitored cortical contractility in embryos expressing either a GFP-tagged form of a wild-type [GFP-AIR-1] or a kinase-dead mutant of AIR-1 [GFP-AIR-1^T201A^] in the absence of the endogenous AIR-1. Notably, we found that GFP-AIR-1 expressing embryos fully rescue the excess cortical contractility phenotype observed upon loss of endogenous protein, which is not the case in the embryos expressing GFP-AIR-1^T201A^(compare Supplementary Figure S1F with S1E and S1G). Overall, this data suggests that AIR-1 kinase function is crucial for proper cortical contractility in the one-cell embryo and PPH-6 phosphatase may modulate this activity (see the Discussion).

**Figure 1:**
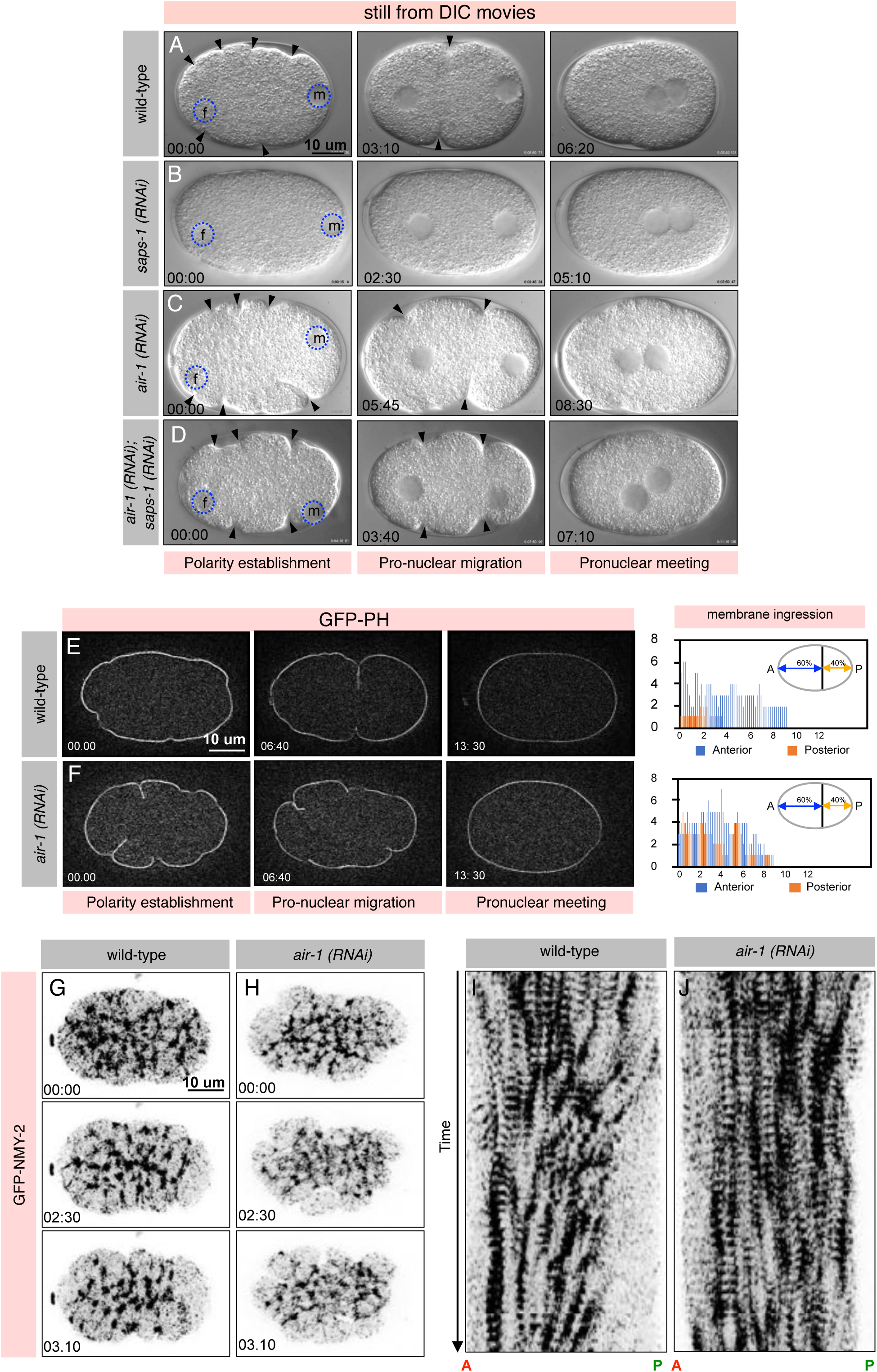
Aurora A kinase (AIR-1) depletion cause excess cortical contractility in the early one-cell stage embryos. (A-D) Images from time-lapse Differential Interference Contrast (DIC) microscopy of *C. elegans* early embryos in wild-type (A), *saps-1(RNAi)* (B), *air-1(RNAi)* (C) or *air-1 (RNAi); saps-1 (RNAi)* (D). See also corresponding Supplementary Movies S1-S4. In this and subsequent images, embryos are approximately 50 μm in length, and the posterior is to the right. Blue dashed circles highlight the male and female pronucleus, and black arrowhead represents the cortical ruffles. In this and other DIC images, if not specified, time is shown in minutes standardize by the size of the male pronucleus and appearance of the posterior smoothening with t=0 corresponding to the size equivalent of 5-6 μm of the male pronucleus (see Materials and Methods for detail). Note excess cortical contractility in the *air-1 (RNAi)* embryo either alone or together with *saps-1 (RNAi)*, in contrast to the wild-type embryo. Also, note significantly diminished cortical contractility in *saps-1 (RNAi)* embryo. More than 50 embryos were analyzed in each condition, and the images from the representative embryos are shown here. (E, F) Images from time-lapse confocal microscopy of embryos expressing GFP-PLCδPH in wild-type (E) and in *air-1 (RNAi)* (F) at the various stages in early embryos. See also corresponding Supplementary Movies S5 and S6. Time is represented in minutes, with t=0 corresponding to the frame from where the imaging was initiated i.e. that is approximately the time of polarity establishment just a few minutes after meiosis II completion. Note excess membrane ruffles in the *air-1(RNAi)* embryo in comparison with the wild-type embryo. Quantification on the right represent membrane ingression in the 60% of the anterior (A) region (shown by blue) and the 40% of the posterior (P) region of the embryo (shown in orange), please see Materials and Methods for the detail of this analysis. Note that in the wild-type embryos the posterior ingression diminishes very early and are absent at the time of pronuclear migration. However, such ingressions stay for a much longer duration and disappear together with the anterior ingression in *air-1 (RNAi)* condition. 10 embryos were analyzed for each condition and representative embryos are shown. (G-J) Cortical images (G, H) and the Kymograph analysis (I, J; see Materials and Methods) from time-lapse confocal microscopy of embryos expressing GFP-NMY-2 in wild-type (G, I) and in *air-1 (RNAi)* (H, J). See also corresponding Supplementary Movies S7 and S8. Time is represented in minutes, with t=0 corresponding to the frame from where the imaging was initiated i.e. that is approximately the time of polarity establishment just a few minutes after meiosis II completion. Note that in the wild-type embryo the GFP-NMY-2 foci move from the posterior (P) to the anterior surface (A) which is not the case in *air-1 (RNAi)* embryos at early time points. More than 10 embryos were recorded, and the represented embryos are shown.

Newly fertilized *C. elegans* zygotes show uniform actomyosin-based cortical contractions that cease at the posterior cortex at the time of polarity establishment (Munro et al., 2004; reviewed in Cowan and Hyman, 2007; Motegi and Seydoux, 2013; Rose and Gönczy, 2014). Therefore, we quantitatively measured cortical contractility in term of membrane ingression (see Figure legends for 1E, 1F and materials and methods for details) in wild-type and *air-1 (RNAi)* embryos expressing the plasma membrane marker GFP-PH. In wild-type embryos, cortical ingressions were uniform at the entire embryo surface before polarity establishment phase, and such ingression ceases on the posterior cortex at the onset of polarity establishment (Figure 1E; Supplementary Movie S5). On the contrary, embryos depleted of AIR-1 show robust cortical ingression at the posterior membrane that stays for a prolonged duration and disappears along with the anterior ingression at the time of the pronuclear meeting (Figure 1F; Supplementary Movie S6).

Next, we examined whether the excess contractility upon AIR-1 loss correlates with the change in the cortical localization of the GFP-NMY-2. Intriguingly, we uncovered that in both wild-type and *air-1(RNAi)* embryos GFP-NMY-2 patches appear similar, however, in comparison with wild-type embryos, *air-1 (RNAi)* embryos do not disassemble GFP-NMY-2 patches at the posterior cortex at the onset of polarity establishment (compare Figure 1H and 1J with 1G and II; Supplementary Movies S7 and S8). To characterize cortical flows further, we conducted particle image velocimetry (PIV) analysis of the wild-type and *air-1 (RNAi)* embryos before and during first 180 s of the GFP-NMY-2 movies (Supplementary Figure S1H-S1K). These analyses clearly indicate that the cortical flows are highly reduced upon AIR-1 loss in comparison with the wild-type embryos. Altogether, this data suggests that uniform ingression-regression on the entire embryo surface upon AIR-1 loss stems from the local activity of actomyosin and the lack of strong cortical flow that is observed in otherwise wild-type embryos.

### AIR-1 ensures single polarity axis in the one-cell embryos

What are the consequences of excess contractility during early cell cycle stages? Using fixed immunostained embryos, AIR-1 depletion is implicated in the impairment in the localization of anterior PAR and, posterior PAR [referred to as aPAR and pPAR respectively], as well as P-granules during late mitotic stages (Schumacher et al., 1998; Noatynska et al., 2010). However, the potential reason by which AIR-1 depletion influences PAR localization remained elusive. Given that impairment in cortical contractility can affect A-P polarity in the one-cell embryo (Motegi et al., 2006; Schonegg and Hyman, 2006; reviewed in Munro, 2006; Cowan and Hyman, 2007) and the timing of excess contractility in *air-1(RNAi)* embryos coincides with the polarity establishment phase, we sought to assess the impact of AIR-1 loss on pPAR using a *C. elegans* strain where the endogenous copy of PAR-2 is tagged with mCherry using CRISPR/Cas9 [mCherry-PAR-2]. Notably, in contrast to the control cells where PAR-2 showed stereotype localization at the posterior cell cortex at the onset of polarity set-up, mCherry-PAR-2 signal appeared at the anterior as well as the posterior cell cortex (Figure 2A-2D and 2H; Supplementary Movies S9 and S10). Similar data are obtained in transgenic strain ectopically expressing GFP-PAR-6 and mCherry-PAR-2 (Supplementary Figure S2A-S2B and Supplementary Movies S11 and S12). Also, the timing of two distinct mCherry-PAR-2 domains appearance in *air-1 (RNAi)* embryos is similar to that of wild-type (Figure 2E-2G).

**Figure 2:**
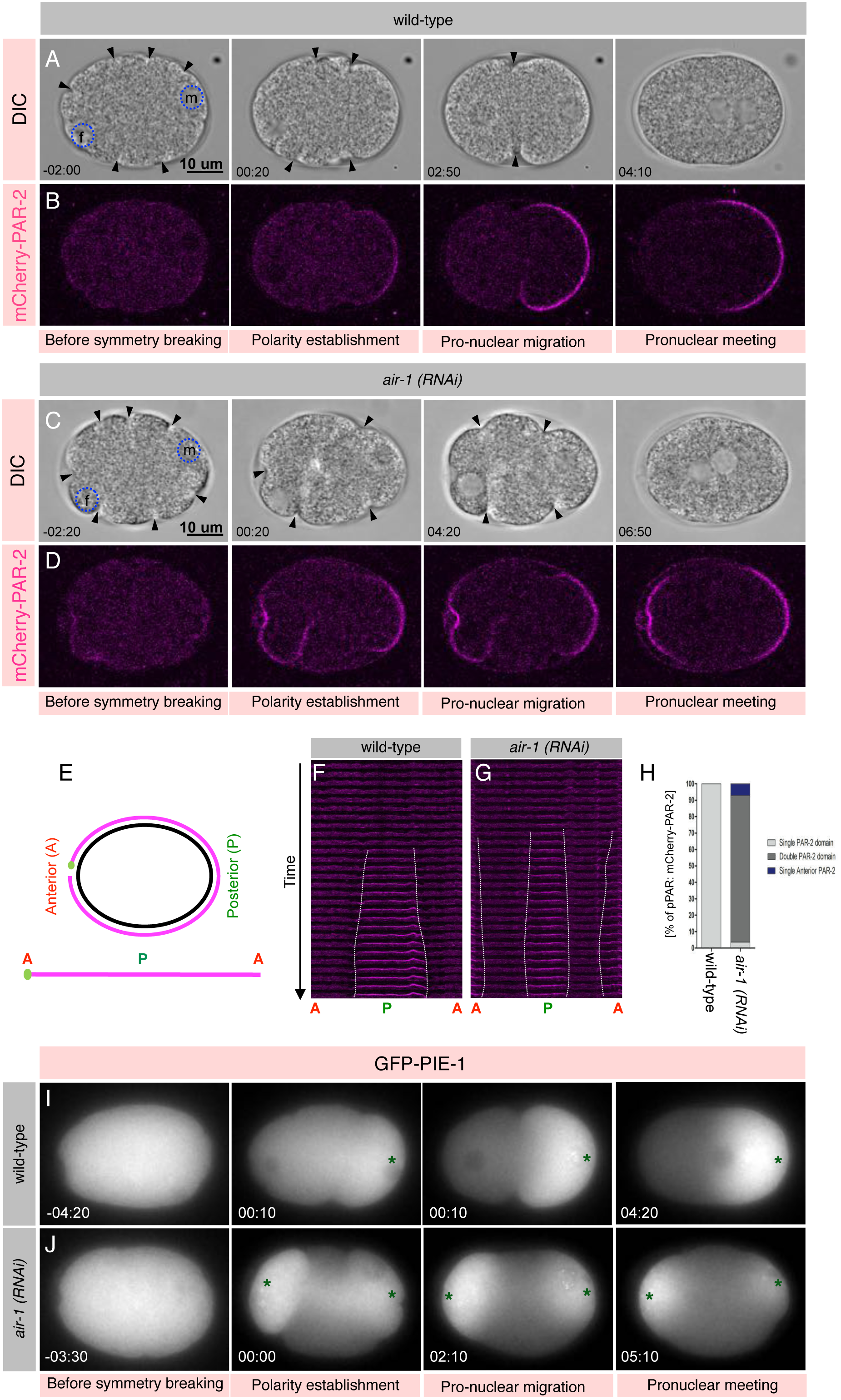
AIR-1 blocks multiple polarization events in the one-cell embryo. (A-D) Images from time-lapse confocal microscopy in combination with DIC microscopy of embryos expressing mCherry-PAR-2 in wild-type (A, B) or in *air-1 (RNAi)* (C, D) at the various stages in the early embryos. See also corresponding Supplementary Movies S9 and S10. mCherry-PAR-2 signal is shown in magenta. Note the presence of excess membrane ruffles in the *air-1(RNAi)* embryo in comparison with the wild-type embryo. Also, note the appearance of two PAR-2 domains at the anterior and posterior cell cortex in the *air-1 (RNAi)* embryo in comparison with the single posterior PAR-2 domain in the wild-type embryos. In this and other images from one-cell embryos expressing either mCherry-PAR-2 or GFP-PAR-2, if not specified, time is shown in minutes standardize by the size of the male pronucleus and the enrichment of the mCherry-PAR-2 signal at the posterior cortex with t=0 corresponding to the size of male pronucleus equivalent of 5-6 μm (see Materials and Methods for detail). More than 50 embryos were recorded and the represented embryos are shown. (E) Schematic of an embryo where the cortical surface (magenta) is flattened from anterior (A) to posterior (P) to anterior (A) to analyze the mCherry-PAR-2 localization in the form of kymograph as shown for F and G. (F, G) Cortical localization of mCherry-PAR-2 signal from the time-lapse confocal images from the 8-10 frames before polarization till the initiation of the pseudocleavage in the one-cell of wild-type embryo or *air-1 (RNAi)* for the analogous timings. Please note that mCherry-PAR-2 signal appears at the A as well as P cortical region in *air-1 (RNAi)* in contrast to the wild-type embryos. Also, note that time timing of PAR-2 domain appearance in *air-1 (RNAi)* embryos are similar to the wild-type embryos. See Material and Methods for the details of this analysis. (H) Quantification of the positioning of the PAR-2 domain appearance in the wild-type and in the embryos depleted of AIR-1. Note the formation of the two PAR-2 axes in the more than 85% of the *air-1 (RNAi)* embryos in contrast to the none in the wild-type. Further note that around 7% of AIR-1 depleted embryos also show the appearance of the single anterior PAR-2 domain. N=50. (I, J) Images from time-lapse epifluorescence microscopy of embryos expressing the germline factor GFP-PIE-1 in wild-type (I) or in *air-1 (RNAi)* (J). See also corresponding Supplementary Movies S13 and S14. GFP-PIE-1 signal is shown in grey. Green asterisk represents the enrichment of PIE-1. Note the presence of two GFP-PIE-1 enriched regions in *air-1 (RNAi)* embryos in comparison with one at the posterior in the wild-type embryos. More than 10 embryos were recorded and the represented embryos are shown.

As another means to check the impact of polarity by AIR-1 depletion, we sought to examine the localization of a germline factor, the PIE-1 protein (Reese et al., 2000) and the P-granule component PGL-1 (Kawasaki et al., 1998) in the early embryos. Interestingly, analogous to the PAR-2 localization, GFP-PIE-1 or GFP-PGL-1 are localized at the anterior and posterior of the one-cell embryo upon AIR-1 depletion in contrast to the wild-type embryos where the signal is mainly restricted to the posterior of the embryo (Figure 2I and 2J; Supplementary Figure S2C and S2D; Supplementary Movies S13-S16).

AIR-1 is enriched at the centrosome as well as on the astral microtubules (Hannak et al., 2001; Kotak et al., 2016). AIR-1 localization at the astral microtubules is mediated by it co-activator TPXL-1 (TPX-2 in human; Supplementary Figure S2E and S2F; Kufer et al., 2002; Hamill et al., 2002; Ozlu et al., 2005; Kotak et al., 2016). Therefore, we sought to examine the contribution of astral microtubule pool of AIR-1 on PAR-2 localization. *tpxl-1(RNAi)* embryos neither show excess contractility nor a promiscuous PAR-2 domain (Supplementary Figure S2G and S2H; Supplementary Movie S17). These data suggest that neither AIR-1 localization at the astral microtubules nor its coactivator TPXL-1 is required for its function in ensuring the formation of single polarity axis at the time of polarity establishment.

### AIR-1’s role in polarity is independent of its role in centrosome maturation

AIR-1 is an essential kinase that is crucial for centrosome maturation and thus bipolar spindle formation (Hannak et al., 2002). Depletion of AIR-1 dramatically impairs α-tubulin fluorescence intensity presumably by impacting the localization of pericentriolar material (PCM) proteins such as γ-tubulin, CeGrip and ZYG-9 (Hannak et al., 2002). Because, microtubules are linked with polarity establishment (Wallenfang and Seydoux, 2000; Tsai and Ahringer, 2007; reviewed in Siegrist and Doe, 2007), we sought to clarify the role of microtubules in the AIR-1-dependent polarity establishment pathway. We reasoned if the appearance of a promiscuous PAR-2 domain in AIR-1 depleted embryos is a result of the diminished number of microtubules, we should mimic this phenotype in a condition that significantly reduces microtubule network in the one-cell embryo. To this end, we performed *tbg-1 (RNAi)* experiment that leads to loss of γ-tubulin as monitored by the collapse of the centrosomes after nuclear envelope break down [NEBD] (Supplementary Movie S18; Hannak et al., 2002). We noticed that loss of γ-tubulin neither impact cortical contractility nor GFP-PAR-2 localization (Figure 3A and 3B). As a more rigorous test to study the role of microtubules in AIR-1 mediated PAR-2 localization, we treated wild-type or AIR-1 depleted early embryos with the microtubule poison Nocodazole and assessed mCherry-PAR-2 localization in such a setting. As described previously (Sonneville and Gönczy, 2004), Nocodazole-mediated de-polymerization of microtubules in the wild-type embryos does not influence PAR-2 localization at the polarity establishment phase. Strikingly, Nocodazole treated *air-1 (RNAi)* embryos still harbour two-PAR-2 domains similar to the embryos solely lacking AIR-1 (Figure 3C-3H; Supplementary Movies S19 and S20). Altogether, this data indicates that AIR-1 function in maintaining single polarity axis is independent of its role in regulating microtubules number at the centrosomes.

**Figure 3:**
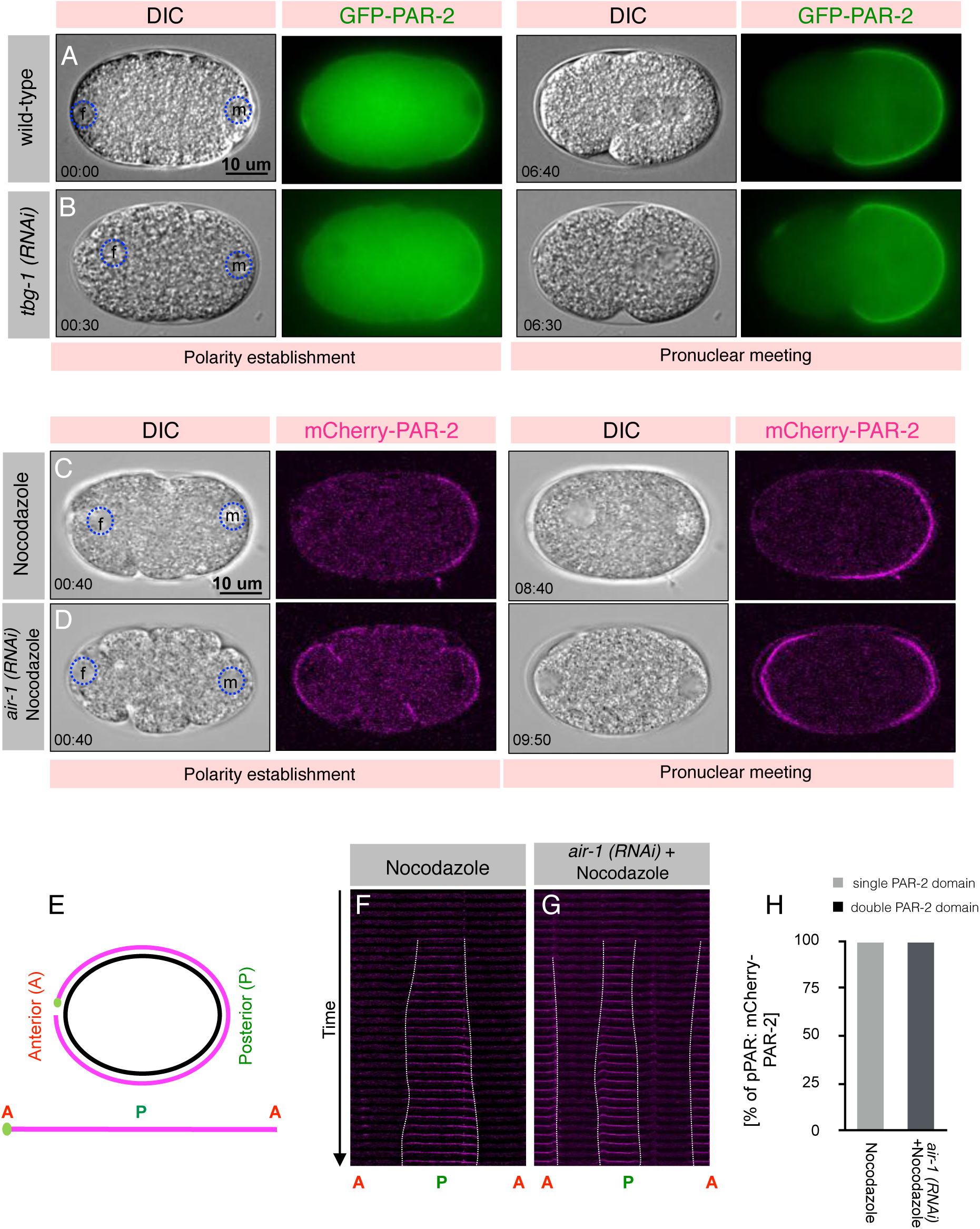
AIR-1 role in maintaining the correct microtubules number is not essential for AIR-1 function in blocking promiscuous pPAR domains. (A, B) Images from time-lapse epifluorescence microscopy in combination with DIC microscopy of embryos expressing GFP-PAR-2 in wild-type (A) or in *tbg-1 (RNAi)* (B). GFP-PAR-2 signal is shown in green. More than 10 embryos were recorded and the represented embryos are shown. Note the *tbg-1 (RNAi)* phenotype in the Supplementary Movie S18. (C, D) Images from time-lapse confocal microscopy in combination with DIC microscopy of wild-type embryos expressing mCherry-PAR-2 and treated with microtubules poison Nocodazole (C) or *air-1 (RNAi)* embryos which are also treated with Nocodazole (D) in early one-cells embryos. See also corresponding Supplementary Movies S19 and S20. mCherry-PAR-2 signal is shown in magenta. Note that the pronuclei do not migrate and stay close to the anterior and posterior cortex at the pronuclear meeting stage because of the absence of the microtubules in Nocodazole-treated cells. More than ten embryos were recorded and the represented embryos are shown. (E) Schematic of an embryo as demonstrated for the Figure 2C for analyzing mCherry-PAR-2 localization over time in the form of kymograph. (F, G) Cortical localization of mCherry-PAR-2 signal from the time-lapse confocal images from 5 frames before polarization till the initiation of the pseudocleavage in wild-type and for the equivalent time in *air-1 (RNAi)* one-cell embryos. Please note that mCherry-PAR-2 signal appears at the A as well as P cortical region in the AIR-1 depleted embryos which are also treated with Nocodazole (G) in comparison with Nocodazole treated embryos where the signal is only present at the posterior cortex (F). Also, note that time timing of PAR-2 domain appearance in *air-1 (RNAi)* plus Nocodazole embryos are similar to the Nocodazole only embryos. See Material and Methods for details of this experimental condition. (H) Quantification of the positioning of the PAR-2 domain appearance in Nocodazole-treated embryos and *air-1 (RNAi)* embryos that are also treated with Nocodazole. Note the formation of the two PAR-2 domain axes in 100% of *air-1 (RNAi)* plus Nocodazole embryos in contrast to the 0% in embryos treated with Nocodazole only. N=10.

### Centrosome position is dispensable for polarity initiation in AIR-1-depleted embryos

In *C. elegans* one-cell embryos polarity establishment depends on centrosomes (O’Connell et al., 2000; Hamill et al., 2002; Cowan and Hyman, 2004). Moreover, it is illustrated that centrosome can initiate polarity from any position within the embryo (Bienkowska and Cowan, 2012). However, centrosome-cortex distance defines the polarity establishment timing, i.e. shorter the distance the faster polarity establishment (Bienkowska and Cowan, 2012). Since AIR-1 is enriched at the centrosomes, the centrosomal pool of AIR-1 could be the key to ensure single polarity axis in one-cell embryos. Therefore, we decided to establish the contribution of centrosome positioning in modulating pPAR localization in embryos also depleted for AIR-1. To test this, we utilized embryos co-expressing GFP-TAC-1 as the centrosomal marker (Bellanger and Gönczy, 2003; Bot N et al., 2003; Srayko et al., 2003) and mCherry-PAR-2 and examined GFP-TAC-1 position and PAR-2 membrane association during polarity establishment. As suggested earlier, we found that in wild-type embryos, mCherry-PAR-2 first appears at the cortex in the close vicinity to the centrosomes (Figure 4A; Supplementary Movie S21; Cowan and Hyman, 2004; Bienkowska and Cowan, 2012), even in embryos which underwent lateral fertilization (Figure 4B; Supplementary Movie S22). To corroborate this more rigorously, we utilized *zyg-12 (RNAi)* condition. ZYG-12 is a Hook family of cytoskeletal linker protein and loss of ZYG-12 perturb attachment of the centrosome to the nucleus and allows centrosomes to float freely in the cytosol in the one-cell embryo (Malone et al., 2003). We observed that in the *zyg-12 (RNAi)* movement of centrosomes towards the anterior drive mCherry-PAR-2 at the anterior instead at the posterior cortex (Figure 4C; Supplementary Movie S23), further strengthening the existing model that centrosome positioning dictates the site for the PAR-2 anchoring. Next, we sought to assess mCherry-PAR-2 localization in AIR-1 depleted embryos where the fertilization is either lateral or centrosome are not associated with the nucleus because of the absence of ZYG-12. Notably, in both scenarios, PAR-2 localizes at the anterior and posterior cortical domains irrespective of the positioning of centrosome (Figure 4D and 4E; Supplementary Movies S24 and S25). These results strongly support the notion that in AIR-1 depleted embryos centrosome lose the capacity to dictate cortical PAR-2 positioning during polarity establishment in the one-cell zygote.

**Figure 4:**
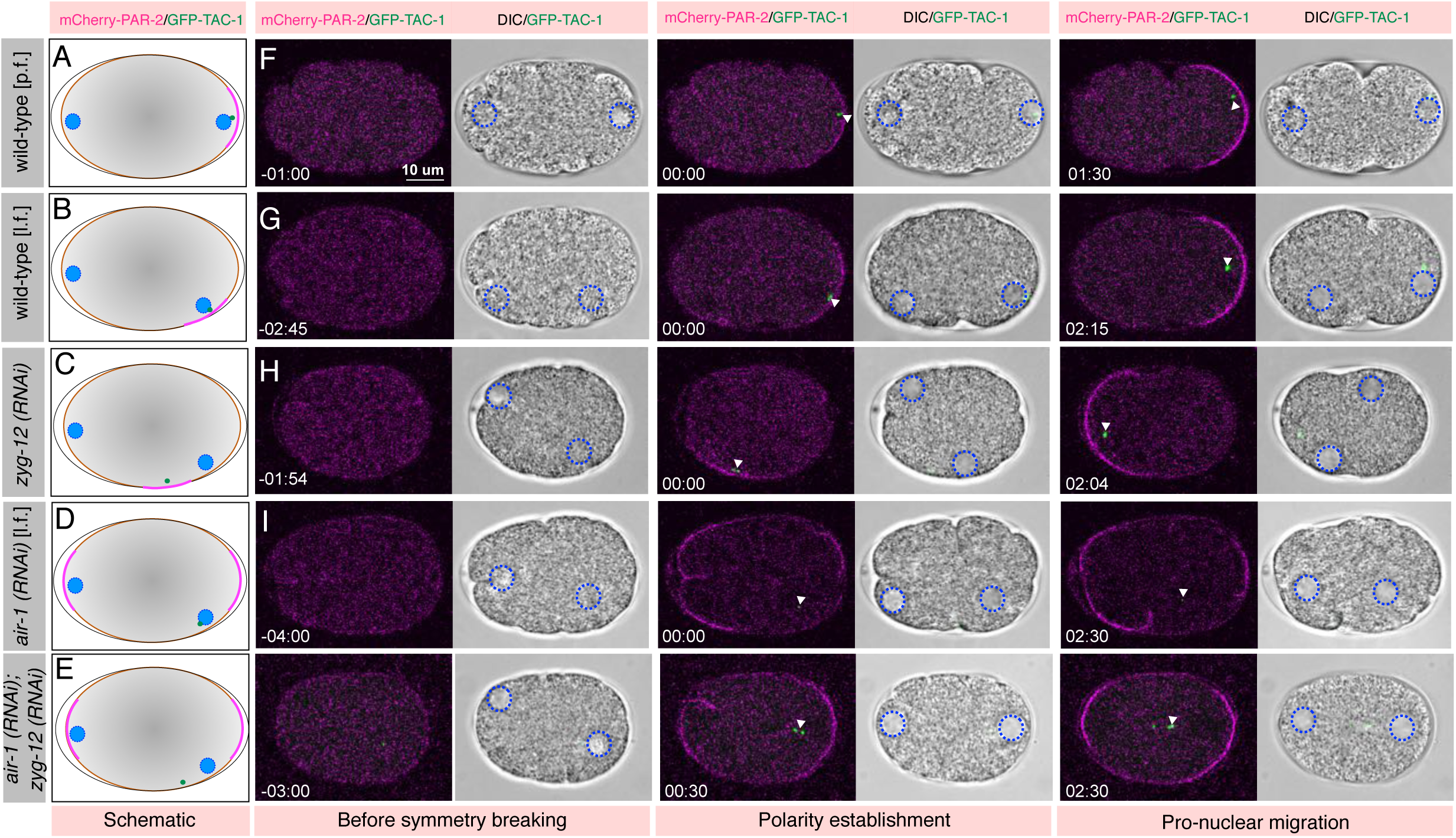
Centrosomes are dispensable in guiding pPAR axis formation in embryos depleted of AIR-1. (A-J) Schematic of the embryos on the left (A-E) represents the various conditions and their corresponding images on the right (F-J) from time-lapse confocal microscopy in combination with DIC microscopy of embryos co-expressing mCherry-PAR-2 and GFP-TAC-1 on the right (F-J). Wild-type embryos underwent posterior fertilization [p.f.] (A, F), wild-type embryos underwent lateral fertilization [l.f.] (B, G), embryos depleted of Hook family of cytoskeletal linker protein ZYG-12 (C, H), *air-1 (RNAi)* embryos underwent lateral fertilization (D, I) or in *air-1 (RNAi) zyg-12 (RNAi)* embryos (E, J) in the early one-cell stages. See also corresponding Supplementary Movies S21-S25. In schematic representations blue circles represent female and male pronuclei, magenta on the cell cortex represent PAR-2 localization and small green circle mark the centrosomes. In confocal images white arrowhead represents the position of the centrosomes. mCherry-PAR-2 signal is shown in magenta. Note that in zyg-12 (RNAi) the centrosomes are not anchored on the nuclear membrane and the position of such unanchored centrosome is sufficient to drive mCherry localization in 4C. Also, note that *air-1 (RNAi)* embryos which either underwent lateral fertilization or in where centrosomes are unable to anchor at the nuclear envelope for instance in *zyg-12 (RNAi)* shows anterior-posterior domain formation irrespective of the position of the centrosomes. More than 7 embryos were recorded for each condition and the represented embryos are shown.

### ECT-2 and its activator NOP-1 acts downstream of AIR-1 in regulating polarity establishment

Thus far our results indicate that AIR-1 at the centrosome could be the key to initiate the symmetry breaking event and thus for the polarity establishment at the embryo posterior. In *C. elegans* one-cell embryo the down-regulation of Rho signalling (Figure 5A) at the posterior cortex at the time of polarity establishment is associated with the membrane exclusion of RhoGEF ECT-2 (Motegi and Sugimoto, 2006). Since AIR-1 loss leads to uniform ingression-regression and non-disassembly of NMY-2 patches (Figure 1) at the posterior cortex, we wondered if ECT-2 or its activator NOP-1 (Tse et al., 2012) acts downstream of AIR-1 in controlling cortical contractility. As expected either, *air-1 (RNAi) ect-2 (RNAi)* or *air-1 (RNAi) nop-1 (RNAi)* embryos display diminished cortical contractility in contrast to *air-1 (RNAi)* (Figure 5B-5E; Supplementary Figure S3A and S3B). Next, we analyzed the impact of ECT-2 or NOP-1 loss on the multiple PAR-2 domains that occur in *air-1 (RNAi)* embryos. Importantly, the occurrence of promiscuous PAR-2 domain upon AIR-1 depletion is completely absent in *air-1 (RNAi) ect-2 (RNAi)* or *air-1 (RNAi) nop-1 (RNAi)* (Figure 5F-5I; Supplementary Figure S3C and S3D; Supplementary Movies S26-S29). However, the posterior PAR-2 domain appearance in *ect-2 (RNAi)* or *air-1 (RNAi); ect-2 (RNAi)* embryos is significantly delayed (Supplementary Figure S3E), and probably in such a condition PAR-2 posterior localization is dependent on the microtubule-directed pathway that kicks in significantly later and polarizes the embryos defective for Rho activation (Zonies et al., 2010; Motegi et al., 2011; Tse et al., 2012; see the Discussion).

**Figure 5:**
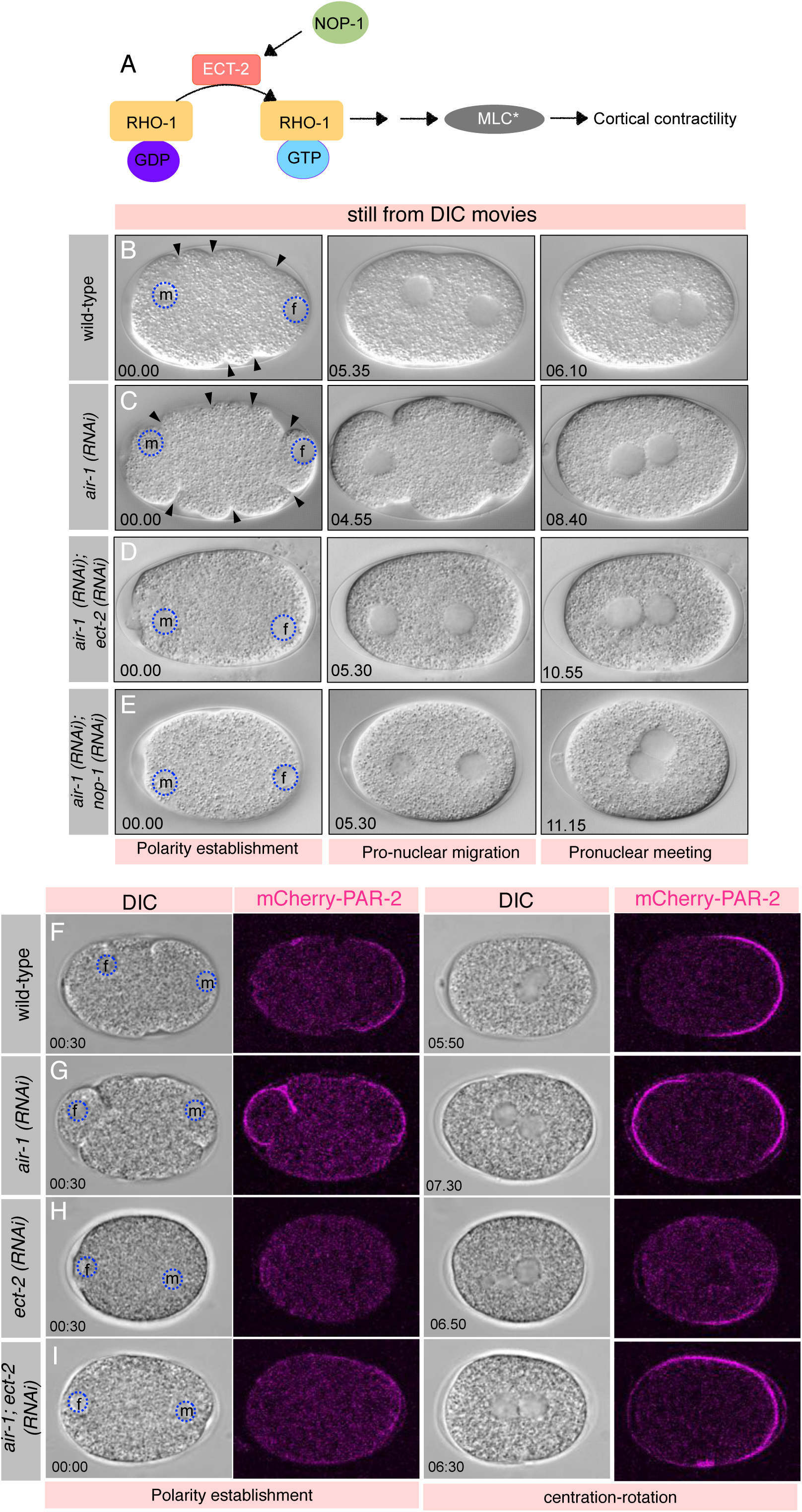
ECT-2 and its activator NOP-1 act downstream of AIR-1. (A) Schematic representation of the pathway regulating cortical contractility by regulating myosin activity in the one-cell embryo. (B-E) Images from time-lapse DIC microscopy of early embryos in wild-type (B), *air-1(RNAi)* (C), *air-1 (RNAi); ect-2 (RNAi)* (D) and *air-1 (RNAi); nop-1 (RNAi)* (E). Note excess cortical contractility that is seen in the *air-1(RNAi)* embryo is diminished in *air-1 (RNAi); ect-2 (RNAi)* or *air-1 (RNAi); nop-1 (RNAi)* embryos. Time is shown in minutes which is solely based on the size of the male pronucleus because of the absence of cortical contractility in *ect-2 (RNAi)* or *nop-1 (RNAi)*. t=0 corresponding to the size equivalent of 5-6 μm. Ten embryos were analyzed in each condition and represented embryos are shown. (F-I) Images from time-lapse confocal microscopy in combination with DIC microscopy of embryos expressing mCherry-PAR-2 in wild-type (F), *air-1 (RNAi)* (G), *ect-2 (RNAi)* (H) and *air-1 (RNAi) ect-2 (RNAi)* (I) at various stages as mentioned. See also corresponding Supplementary Movies S26-S29. mCherry-PAR-2 signal is shown in magenta. Note the appearance of an extra PAR-2 domain at the anterior in *air-1 (RNAi)* is lost in *air-1(RNAi); ect-2 (RNAi)*. Also note that in *ect-2 (RNAi)* or in *air-1(RNAi) ect-2 (RNAi)* shows the appearance of the posterior PAR-2 domain that is significantly delayed than wild-type as reported earlier for ECT-2 depletion (see Supplementary Figure S3E and Tse et al., 2012). More than 10 embryos were recorded for each condition except *air-1 (RNAi); ect-2 (RNAi)* where n=5 and the represented embryo are shown.

### AIR-1 delocalizes ECT-2 at the posterior cortex at the time of polarity establishment

Since the loss of ECT-2 localization a the posterior cortex set-up anisotropy in the one-cell embryos to establish aPAR and pPAR domains (reviewed in Hoege and Hyman, 2013), we decided to check ECT-2 localization in embryos lacking AIR-1. To this end we utilized embryos co-expressing GFP-ECT-2 and mCherry-PAR-2 and analyzed the cortical localization of these proteins in the one-cell early embryos. In wild-type embryo as reported earlier, we noticed the exclusion of GFP-ECT-2 at the posterior cortex (Figure 6A, 6E and 6J; Motegi et al., 2006) and this coincides with the localization of mCherry-PAR-2 at the posterior cortex (Figure 6B, 6F and 6K; Supplementary Movie S30). Interestingly, in AIR-1 depleted embryos GFP-ECT-2 remains associated with the posterior cortex and two mCherry-PAR-2 domains are formed (Figure 6C, 6D, 6G, 6H, 6L and 6M; Supplementary Movie S31). Taken together, our data support the hypothesis that centrosome localized AIR-1 gradient sets formation of a single polarity axis by delocalizing ECT-2 from the posterior cortex and in the absence of AIR-1 relatively uniform cortical contractility driven by non-exclusion of the RhoGEF ECT-2 from the posterior cortex causes the formation of multiple polarity axes (see the Discussion).

**Figure 6:**
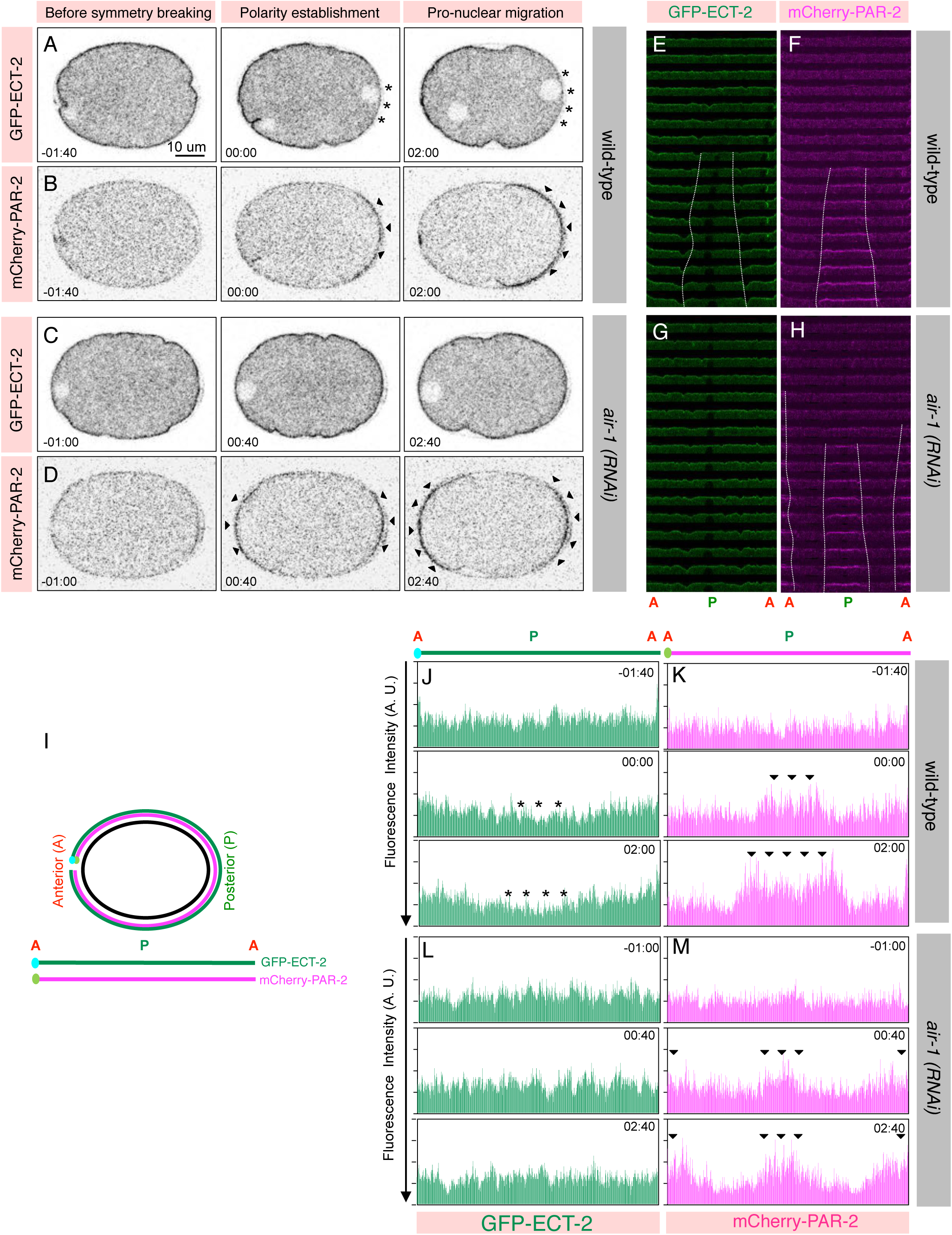
AIR-1 delocalizes ECT-2 at the posterior cortex at the time corresponding to polarity establishment. (A-D) Inverted contrast images from time-lapse confocal microscopy of one-cell embryos co-expressing mCherry-PAR-2 and GFP-ECT-2 in wild-type (A, B) or *air-1 (RNAi)* (C, D) at the various stages as mentioned. See also corresponding Supplementary Movies S30 and S31. mCherry-PAR-2 and GFP-ECT-2 signal is shown in grey. Note that GFP-ECT-2 delocalize at the posterior cortex at the time of symmetry breaking in the one-cell embryo and a concomitantly mCherry-PAR-2 signal appears at the posterior cortex. Also, note that in the *air-1 (RNAi)* two PAR-2 axis are established and ECT-2 do not delocalize from the posterior cortex. More than 5 embryos were recorded and the represented embryos are shown. (E-H) Cortical localization of GFP-ECT-2 and mCherry-PAR-2 signal from the time-lapse confocal images from the 8 frames before polarization till the initiation of the pseudocleavage in the one-cell embryo as shown for 2E-2G. Please note that GFP-ECT-2 exclusion coincide with the appearance of mCherry-PAR-2 at the posterior cell cortex in wild-type embryos. Also note that GFP-ECT-2 signal is not diminished from the posterior cortex in *air-1 (RNAi)* and mCherry-PAR-2 signal appears at the A as well as P cortical surface. (I) Schematic of an embryo where the cortical surface (green for GFP-ECT-2 or magenta for mCherry-PAR-2) is flattened from anterior (A) to posterior (P) to anterior (A) to analyze the fluorescence cortical intensity overtime. (J-M) Line scan analysis of the fluorescence intensities in arbitrary units [A.U.] for the cortical GFP-ECT-2 (in green) or mCherry-PAR-2 (in magenta) for the images shown in A-D in wild-type (J, K) or *air-1 (RNAi)* (L, M). Asterisks represent the cortical region that is depleted of GFP-ECT-2 and black arrowheads represent the cortical region where mCherry-PAR-2 signal appears in the embryos. Note the appearance of mCherry-PAR-2 coincide with the loss of cortical GFP-ECT-2.

## Discussion

The proper establishment of polarity axis during various stages of development is vital for regulating multiple events such as cell division, cell migration and cell signalling. Across multiple cell types, distinct signals are at play to trigger polarity establishment. For instance, epithelial cells polarize in response to the extrinsic cues from their neighbours, mating yeast cells polarize in response to the signals from each other and *Caenorhabditis elegans* one-cell stage embryos polarize by the sperm entry just after fertilization (reviewed in Chant, 1999; Nelson, 2003; Pellettieri and Seydoux, 2002; St Johnston, 2018). In the one-cell of *C. elegans* zygote, the cortical actomyosin network is considered to act as a significant driving force for the asymmetric distribution of the molecules that are responsible for the polarity establishments (Munro et al., 2004; reviewed in Munro and Bowerman, 2009). In this study, we demonstrate a novel function of AIR-1 (Aurora A) kinase in modulating actomyosin network through the RhoGEF ECT-2, and we uncover that this action of AIR-1 is critical for ensuring the establishment of single polarity axis.

### AIR-1 is crucial for regulating proper cortical contractility in the one-cell embryo

Our results demonstrate that severe depletion of AIR-1 cause much pronounced cortical contractions and this correlates well with the timing of polarity establishment (Figure 7A and 7B). Impact of AIR-1 loss on cortical contractility is antagonistic to the loss of PP6 phosphatase regulatory subunit SAPS-1. Further, by conducting epistasis experiments, we revealed that AIR-1 acts downstream of SAPS-1 in controlling cortical contractility. In human cells, PP6 acts as a major T-loop phosphatase for Aurora A (Zeng et al., 2010), and thus it may well be that the analogous situation exists in *C. elegans* zygote whereby PPH-6/SAPS-1 phosphatase complex buffer AIR-1 activity to modulate proper cortical contractility. Notably, in the line of this hypothesis, we uncovered that AIR-1 kinase activity is crucial for this function and also discovered that TPXL-1 (Ozlü et al., 2005; Mangal et al., 2018), that is required to control a subset of AIR-1 activity is not essential for AIR-1-mediated cortical contractility.

**Figure 7:**
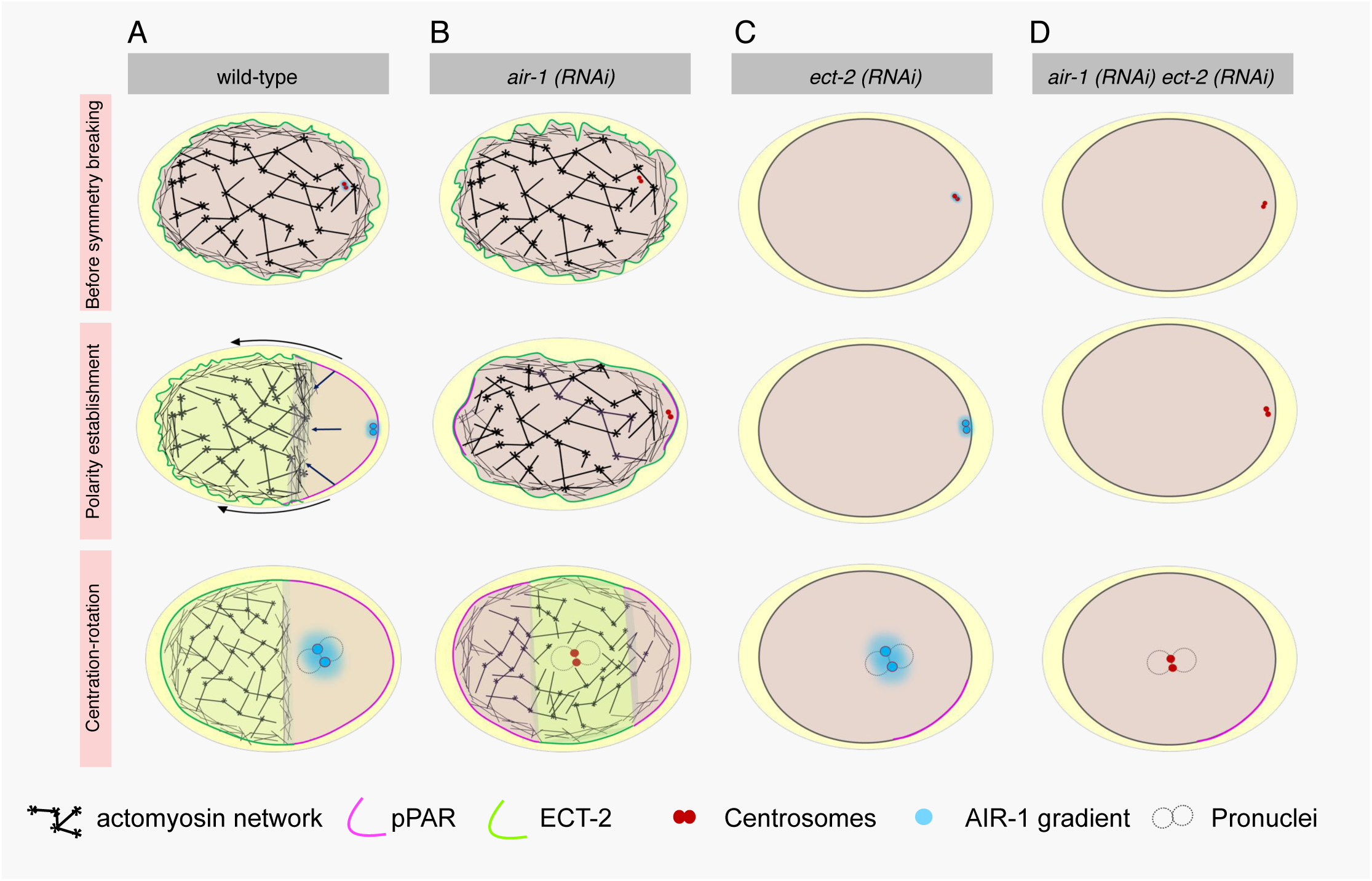
AIR-1 is essential in ensuring single pPAR-2 axis. (A-D) Working model for the function of AIR-1 in regulating single polarity axis at the time of polarity establishment in the one-cell embryo. In the wild-type, before symmetry breaking event, the entire embryo surface is undergoing contraction-relaxation and the RhoGEF, ECT-2 (in green) is localized at the cell membrane and PAR-2 (in magenta) is restricted to the cytoplasm (A). At the onset of polarity establishment, relaxation of the posterior cortex presumably by the exclusion of ECT-2 from the surface close to the AIR-1 centrosomal gradient (in cyan) cause PAR-2 to now localize at the posterior domain and thus enabling polarity establishment. At the centration-rotation stage, a well-defined PAR-2 domain is formed and ECT-2 is mainly restricted to the anterior embryo surface. In *air-1 (RNAi)* embryos an excess cortical contractility is present because of the presence of ECT-2 at the entire surface and such contractility is not abolished at the time of polarity establishment and this cause the occurrence of promiscuous PAR-2 domain at the anterior and at the posterior cortex (B). In contrast in *ect-2 (RNAi)* or on *air-1 (RNAi) ect-2 (RNAi)* embryos contractility is absent and these embryos are characterized by the appearance of the PAR-2 domain that is significantly delayed than the wild-type embryos (C, D). Centrosomes are shown in red.

How AIR-1 influence cortical contractility in the one-cell embryo? AIR-1 is enriched at the centrosome and AIR-1 depletion cause excess actomyosin-based contractility. This observation suggests that in the wild-type embryos AIR-1 gradient at the centrosome might be the key to assist in symmetry breaking event in the vicinity of centrosome for the accurate polarity establishment. In a line of this, we discovered that AIR-1 depletion in the one-cell embryo at the time of polarity set-up blocks the RhoGEF ECT-2 clearance from the cell cortex and concomitantly promotes the formation of spontaneous pPAR domains on the anterior as well as the posterior cell cortex (Figure 7C and 7D). Non-exclusion of ECT-2 from the posterior cell cortex in *air-1 (RNAi)* appears to be the cause for excess contractility.

Based on the above observations we propose that centrosome localized AIR-1 gradient could serve as a diffusive signal that determines the positioning of the single pPAR axis formation (Figure 7). This hypothesis is in line with the previous reports whereby loss of SPD-5, a large coiled-coil protein, essential for AIR-1 recruitment at the centrosomes cause formation of more than one PAR-2 domains in significant number of embryos (Hamill et al., 2002; Tsai and Ahringer, 2007) and also on the findings that the centrosome distance from the closest cell cortex is the primary determining factor for the timing/axis of polarity establishment (Bienkowska and Cowan, 2012). Importantly, our data further indicate that in the absence of AIR-1, the centrosome requirement and their positioning is expendable in guiding the establishment of accurate polarity axis.

### ECT-2: an evolutionarily conserved RhoGEF at the front line for the proper polarity axis formation

ECT-2 is evolutionarily conserved RhoGEF that play key role in polarity establishment (Liu et al., 2004; Motegi and Sugimoto, 2006; Liu at al., 2006; Rosa et al., 2015), cell rounding (Matthews et al., 2012) and for the formation of the cleavage furrow (Prokopenko et al., 1999; Tatsumoto et al., 1999; Dechant and Glotzer, 2003; Yuce et al., 2005). In mammalian cells, the C-terminal of Ect2 possesses PH and PBC domains that are responsible for its interaction with membrane phosphoinositides (Chalamalasetty et al., 2006; Su et al., 2011). There ectopic localization of Ect2 at the plasma membrane is associated with the excess membrane ruffling at the time of furrow formation by regulating the cortical actomyosin activity (Su et al., 2011). Our data reveal that in *C. elegans* one-cell embryo ECT-2 uniformly decorates the entire embryo surface upon AIR-1 depletion and thus it is worth assuming that during polarity establishment ECT-2 similar to its mammalian counterpart rely on the phosphoinositides-based membrane lipids and AIR-1 directly or indirectly helps in changing the lipid binding property of ECT-2 and therefore helps in breaking symmetry. Future work is required to investigate whether ECT-2 could acts as a direct target of AIR-1 in this paradigm.

### Microtubules are not essential for proper polarity establishment in the context of AIR-1

The role of microtubules in the anterior-posterior axis formation is well documented in several model organisms including in the *C. elegans* one-cell embryo (Tsai and Ahringer, 2007; Wallenfang and Seydoux, 2000; reviewed in Siegrist and Doe, 2007). The mutants which are defective in the centrosome maturation [i.e. spd-2 and spd-5] show delay in the microtubules nucleation and thus are one of the bases of the proposal that microtubules might be needed for proper polarity establishment (O‵Connell et al., 2000; Hamill et al., 2002). Keeping this in mind, we have tested whether AIR-1 influences polarity establishment via its proposed function related to centrosome maturation and thus affecting the total number of microtubules (Hannak et al., 2001). Intriguingly, embryos depleted of γ-tubulin that also perturb the centrosome maturation and microtubules nucleation does not show polarity establishment defects in contrast to the AIR-1 loss. Also, Nocodazole-mediated microtubules de-polymerization in the one-cell embryo does not impact polarity establishment as measured by the PAR-2 cortical occupancy either in the wild-type or *air-1 (RNAi)* embryos. In particular, we have unravelled that the position of centrosome and its distance from the cortex become obsolete in guiding the polarity axis in *air-1 (RNAi)* background through the centrosomes are still capable of capable of emanating microtubules albeit less efficiently (Hannak et al., 2001). This observation further suggests that microtubules function of the centrosome is not required when AIR-1 is depleted.

### Loss of AIR-1 cause formation of promiscuous pPAR domains

Loss of AIR-1 lead to the formation of two pPAR axis at the time of polarity set-up. This outcome was surprizing to us, if the centrosomal pool of AIR-1 is the positive regulator of creating anisotropy and thereby creating only single PAR-2 in the wild-type embryos then the loss of AIR-1 should not establish polarity at all? Since two pPAR axis that forms upon AIR-1 depletion can be rescued by simultaneously depleting either ECT-2 or NOP-1, it appears that the cortical contractions on the entire embryo surface upon *air-1 (RNAi)* are the sole reason for the appearance of the two PAR-2 domains. In fact, co-depletion of AIR-1 and ECT-2 significantly delay PAR-2 domain formation in contrast to the wild-type embryos (Figure 6 and Supplementary Figure S3E). In such embryos cortical PAR-2 only appears at the time of pronuclear meeting probably because of the existence of microtubule-directed pathway that can polarize embryos in the absence of Rho-mediated cortical contractility (Figure 7C and 7D; Zonies et al., 2010; Motegi et al., 2011; Tse et al., 2012). What is more puzzling however that what makes the position of such domains mostly invariable and these are primarily restricted only to the anterior and posterior polar cortical regions? Since the cell cortex is characterized by visco-elastic property where the dynamic nodes of myosin hold the underlying actin network (Munro et al., 2004; reviewed in Hoege and Hyman, 2013), it is possible that because of curvature at the very anterior and posterior domain of embryo makes these nodes less connected in comparison with the rest of embryo and such locations are more amenable for symmetry breaking. Another possibility could well be that the curvature could alter the orientation of the actin cytoskeleton such that actin is more aligned in the lateral region of the embryo than at the poles in the *air-1 (RNAi)* embryos. In future, it would be interesting to dissect if the physical-basis of the embryo shape are at play to break symmetry in the *air-1 (RNAi)* background.

In mutants arrested in meiosis, the meiotic spindle can guide the formation of the PAR-2 domain at the anterior cortex (Wallenfang and Seydoux, 2000).Therefore, one can assume that in the wild-type embryos AIR-1 blocks the ability of the meiotic spindle to form the anterior PAR-2 and loss of AIR-1 lead to an ectopic PAR-2 domain establishment at the anterior cortex. This notion appears to be unlikely because (1) we did not observe any significant change in the timing of meiosis completion in the absence of AIR-1 in contrast to the wild-type embryos (data not shown and Schumacher et al., 1998), suggesting that the appearance of PAR-2 at the anterior cortex is not because of meiotic error in *air-1 (RNAi)*. (2) Also, if AIR-1 but not cortical contractility is responsible for the formation of the ectopic PAR-2 domain at the anterior then we do not expect to see a rescue in the *air-1 (RNAi); ect-2 (RNAi)* background. We suspect that the pPAR domain appearance in mutants arrested in meiosis could be related to the overtime protection of PAR-2 from aPKC-mediated phosphorylation (Motegi et al., 2011) by the meiotic spindle thus allowing PAR-2 to access the nearest anterior cortex in such conditions.

Overall, our study addresses an important question and shed light on the long-standing puzzle by illustrating that AIR-1 could act as an elusive polarizing signal at the centrosome that initiates polarity establishment via its impact on actomyosin-based cortical contractility. Since Aurora A is an evolutionarily conserved gene, it is likely that the effect of Aurora A on actomyosin-mediated cortical contractions is a general paradigm that is relevant to various biological processes beyond polarity establishment.

## Materials and Methods

### Strains, RNAi and drug treatment

*C. elegans* wild-type (N2) as well as transgenic lines expressing mCherry-PAR-2 (KK1254), RNAi-resistant GFP-AIR-1R or a kinase-dead mutant equivalent (GFP-AIR-1R-T201A) (Toya et al., 2011), GFP-NMY-2 (Munro et al., 2004), GFP-PH (Audhya et al., 2005); GFP-PAR-2; mCherry-PAR-6 (GZ869), GFP-TAC-1 (Bellanger and Gönczy, 2003); GFP-ECT-2 (Motegi and Sugimoto, 2006), GFP-PIE-1 (JH2015) and GFP-PGL-1 (SS747) were maintained at 24^°^C. Bacterial RNAi feeding strains for *saps-1, air-1, ect-2, nop-1, tbg-1, zyg-12* were obtained either from the *C. elegans* ORFeome RNAi library or Source BioScience (Kamath et al., 2003). *tpxl-1 (RNAi)* construct has been described earlier (Ozlu et al., 2005) as the air-1^N^(RNAi) feeding strain (Toya et al., 2011; Kotak et al., 2016). RNAi against *saps-1, air-1, air-1*^*N*^*(RNAi), saps-1 air-1, tbg-1, tpxl-1, zyg-12, ect-2, air-1(RNAi) ect-2 (RNAi), air-1 (RNAi) nop-1 (RNAi)* was performed by feeding animals starting at the L2 or L3 stage with bacteria expressing the corresponding dsRNAs at 20^°^C or 24^°^C for 30-40 hours before analysis. While performing double depletion using RNAi, the single depletion was performed side by side to observe the single RNAi-mediated phenotype. Also, in cases where RNAi-mediated double depletion mask the phenotype/s of any individual RNAi such as in *saps-1 (RNAi) air-1 (RNAi)* where the phenotype stem from the AIR-1 loss mask the *saps-1 (RNAi)* phenotype, the standardization was done by conducting immunostaining analysis using SAPS-1 antibodies to validate the efficient depletion (not shown).

Microtubules poison Nocodazole treatment was performed as described (Bienkowska and Cowan, 2012). In brief, worms were dissected in Nocodazole (10 μg/ml; Sigma Aldrich, M1404) containing egg buffer (118 mM NaCl, 40 mM KCl, 3.4 mM MgCl2, 3.4 mM CaCl2 and 5 mM HEPES pH 7.4; also see Boyd, 1996) and drug could enter in the embryos because of the permeability of the eggshell during meiosis II (Johnston et al., 2006). The efficiency of Nocodazole was determined by the inability of the pronuclei to migrate.

### Time-lapse microscopy

For most experiments, gravid worms were dissected in M9 or egg buffer and transferred onto a 2% agarose pad containing slides using a mouth pipette. These were then covered with a 20 × 20 mm coverslip. Time-lapse Differential Interference Contrast (DIC) microscopy, dual DIC and confocal microscopy or DIC in combination with fluorescence microscopy were performed on such embryos either on IX53 (Olympus Corporation, Japan) with Qimaging Micropublisher 5.0 Colour CCD Camera (Qimaging, Canada) with 100X 1.4 NA, FV3000 Confocal system with high-sensitivity cooled GaAsP detection unit (Olympus Corporation, Japan) using 60X 1.4 NA objective, or IX83 with XM10 cool CCD chip camera (Olympus Corporation, Japan) using 60X 1.4 NA objective. Images were collected at the intervals of 5-20 seconds per frame Movies were subsequently processed using ImageJ, QuickTime and Adobe Photoshop maintaining relative image intensities within a series. Z-stack series were projected as maximum intensity projections for embryos expressing GFP-TAC-1.

### Indirect immunofluorescence

Embryo fixation and staining for indirect immunofluorescence was performed essentially as described (Gönczy et al., 1999), using 1:200 mouse anti-α-tubulin antibodies (DM1A, Sigma), in combination with 1:200 rabbit anti-AIR-1 (Hannak et al., 2001). Embryos were fixed in methanol at –20°C for 30 minutes and incubated with primary antibodies for one hour at room temperature. Secondary antibodies were Alexa488-coupled antimouse and Alexa568-coupled anti-rabbit, both used at 1:500.

Confocal images were acquired on a FV3000 Confocal system with high-sensitivity cooled GaAsP detection unit (Olympus Corporation, Japan) using 60X objective with NA 1.4 oil and processed in ImageJ and Adobe Photoshop, maintaining relative image intensities.

### Data analysis

Fiji (https://fiji.sc/), GraphPad and MATLAB (MathWorks) were used to perform quantitative analysis. For cortical line scans the cortex of the embryo was manually tracked using the Freehand segmented tool on Fiji. Intensity plot of line thickness equivalent of 7 pixels was performed and values were further smoothened using movmean filter on MATLAB. Kymograph analysis of embryos expressing GFP-NMY-2 was performed as described earlier (Munro et al., 2004). In brief, a section of 8-pixel thickness was extracted roughly from the centre of each image from the images coming from the time-lapse recordings. Thereafter, these images were concatenated vertically using MATLAB. For mCherry-PAR-2 and GFP-ECT-2 kymographs, 50 pixels thick section of the cortex was manually tracked and straightened on Fiji. Resulted images of the time-lapse series were concatenated vertically to generate kymograph. Laplace edge detection filter (Cell Dimension Software, Olympus Corporation, Japan) was used to illustrate images from the GFP-PH embryos. On time-lapse images coming from the embryos expressing GFP-PH, the number of ingressions were calculated in the 60% anterior (A) and 40% posterior (P) of the embryos and the ingression were measured on each frame of the recording in such a way that if an ingression is present in two consecutive frames then it is counted for both of the frames.

### Assigning time ‘0’ for embryos expressing GFP-NMY-2 and GFP-PH

For GFP-NMY-2 cortical imaging and GFP-PH imaging, t=0 corresponding to the frame from where the imaging was initiated i.e. that is approximately the time of polarity establishment just a few minutes after meiosis II completion.

### Assigning time ‘0’ for embryos expressing the fluorescently-tagged PAR-2 protein

It is reported previously that at the time of completion of female meiosis the size of the male pronucleus is 3 μm (Bienkowska et al., 2012). In our time-lapse recordings, we uncovered that the size of the male pronucleus is around 5-6 μm at the time of posterior smoothening or symmetry breaking (referred to as polarity establishment). Since male and female pronuclei are next to the posterior and anterior cortex at this time, their position is also considered as a proxy for assigning the timing for the polarity establishment. Because of the lack of cortical flow in embryos depleted for ECT-2 or NOP-1 either alone or in combination with AIR-1, the polarity establishment phase (time ‘0’) is only determined by the size of the pronucleus (5-6 μm). For assigning time ‘0’ in DIC recordings, we analyzed the appearance of the non-contractile posterior domain and also checked the diameter of the male pronucleus that is never beyond 5-6 μm at this point of time.

### Assigning time ‘0’ for embryos expressing GFP-PIE-1

For assigning time ‘0’ in GFP-PIE-1 expressing worms, we analyzed the appearance of the non-contractile posterior domain and also checked the diameter of the male pronucleus that is never beyond 5-6 μm at this time.

### Particle Image Velocimetry (PIV) analysis

Particle Image Velocimetry (PIV) analysis of NMY-2 cortical flow was performed on maximum intensity projections of sequential time-lapse images from GFP-NMY-2 recordings, using a freely available PIV algorithm, PIVlab 1.32 (can be downloaded from http://pivlab.blogspot.com/). A 171 x 85 pixel (35.42 x 17.61 μm) region of interest was applied to standardize cortex area excluding the edges of the embryo. Single-pass PIV with interrogation area of 24×12 pixels with 50% overlap was applied, as described in Nishikawa et al., 2017. Under the available post-processing options, vector validation was done by setting the threshold to stdev=7, and data smoothening was performed. Obtained values of the velocity field were plotted using custom-written MATLAB program, with the help of MATLAB functions obtained from Ronen Zaidel-Bar’s laboratory.

## Acknowledgements

We thank Asako Sugimoto (Tohoku Uni., Tohoku), Pierre Gönczy (EPFL, Lausanne), Nathan Goehring (The Francis Crick Institute, London), Subramaniam, K (IIT, Madras) and Anthony Hyman (MPI, Dresden) for sharing their precious reagents with us. We are thankful to Daniel St Johnston (The Gurdon Institute, Cambridge), Nathan Goehring (The Francis Crick Institute, London), Mark Petronczki (Boehringer Ingelheim Regional Center, Vienna), Sveta Chakrabarti (IISc, Bangalore), Phong Tran (Institut Curie, Paris) and entire Kotak lab for providing us critical comments on the manuscript. We are thankful for Pierre Gönczy for allowing us to conduct some initial DIC work in his lab and Coralie Busso for her help in swiftly arranging all the reagents for shipment at the time of Kotak lab establishment. We greatly appreciate the help from Prerna Sharma and Debasmita Mondal (Department of Physics, IISc) in particle image velocimetry (PIV) analysis of GFP-NMY-2 movies. We thank Ong Hui Ting and Ronen Zaidel-Bar (Sackler Faculty of Medicine, Tel Aviv) for providing us the Matlab code for PIV analysis. We also thank Caenorhabditis Genetics Center (CGC) for providing us all the required worm strains. We are grateful to the DST-FIST, UGC Centre for the Advanced Study, DBT-IISc Partnership Program and IISc for the infrastructure support. This work is supported by the grants from the Wellcome Trust DBT-India Alliance [grant number: IA/I/15/2/502077] to SK. SK is a Wellcome Trust DBT-India Alliance Intermediate Fellow.

## Supplementary Figure legends

**Supplementary Figure 1:**
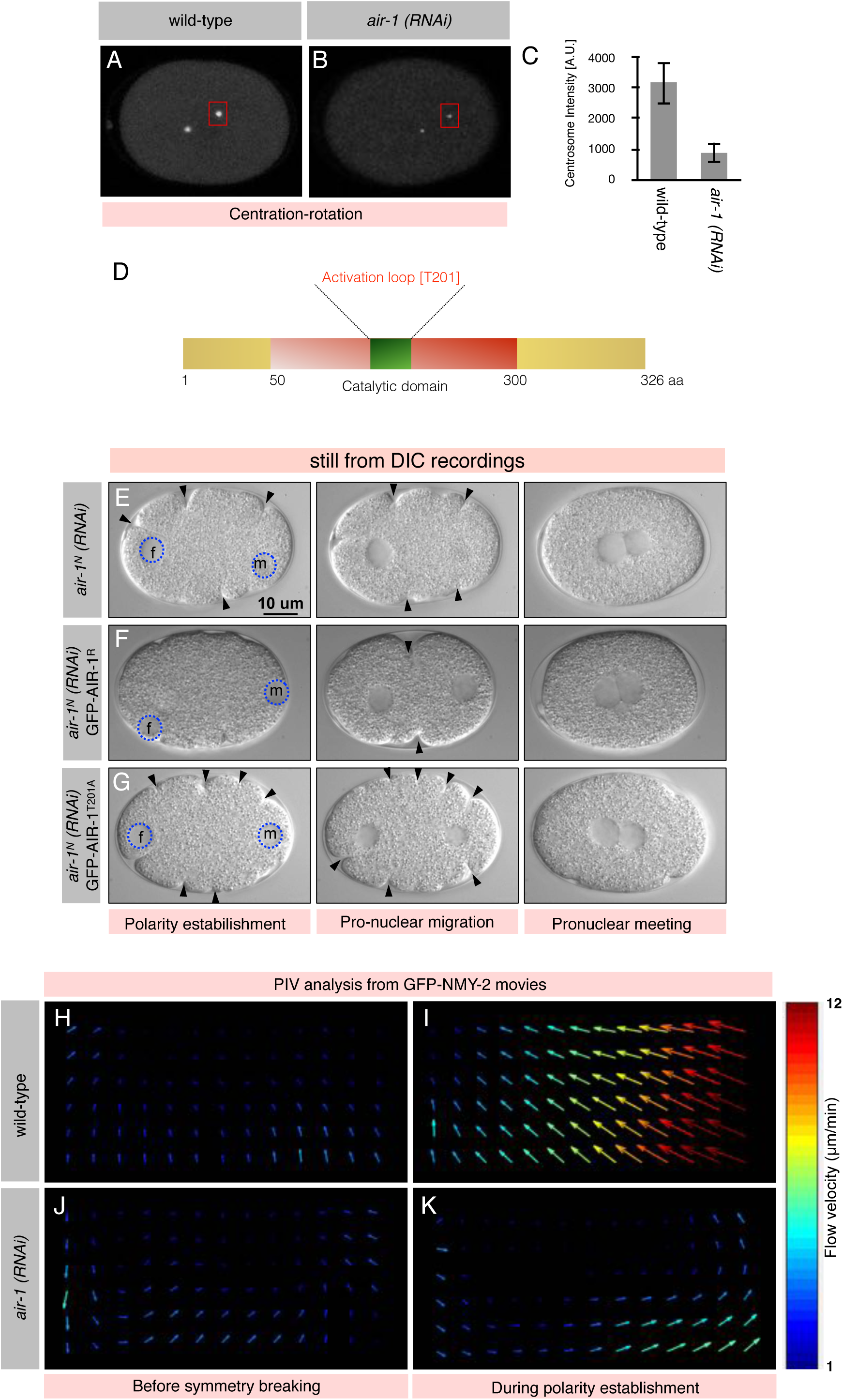
Excess contractility in *air-1(RNAi)* embryos is AIR-1 kinase function dependent. (A-C) Images from time-lapse confocal microscopy from embryos expressing GFP-TAC-1 (shown in grey) as a centrosomal marker in wild-type (A) or *air-1 (RNAi)* condition (B). Centrosome intensity in arbitrary unit [A. U.] was measured (C) by utilizing ten centrosomes from five wild-type and *air-1 (RNAi)* embryos at the centration-rotation stage (see Materials and Methods for details). In this and in subsequent images embryos are approximately 50 μm in length, and posterior is to the right. (D) Schematic of AIR-1 with the position of the catalytic domain (green) with the autophosphorylation site [T201] is indicated. (E-G) Images from time-lapse DIC microscopy of *C. elegans* early embryos in air-1^N^(RNAi) (E), *air-1*^*N*^*(RNAi)* in embryos expressing RNAi resistant form of GFP-AIR-1^R^ (F), or *air-1*^*N*^*(RNAi)* in embryos expressing RNAi resistant form of a kinase-dead [T201A] variant of AIR-1 (GFP-AIR-1^T201A^). Blue dashed circles highlight the male and female pronuclei, and black arrowhead represents the cortical ruffles. Note excess cortical contractility in the *air-1(RNAi)* embryo and in embryos expressing kinase-dead form of AIR-1, but not in the condition that is expressing kinase active form. 10 embryos were analyzed in each condition, and the images from the represented embryos are shown. (H-K) Particle image velocimetry (PIV) analysis of GFP-NMY-2 movies in wild-type (H, I), or *air-1 (RNAi)* (J, K) before and during first 3 min of polarity establishment. In the PIV images, colour arrows indicate direction of the cortical flow and the magnitude of the velocity in um/min.

**Supplementary Figure 2:**
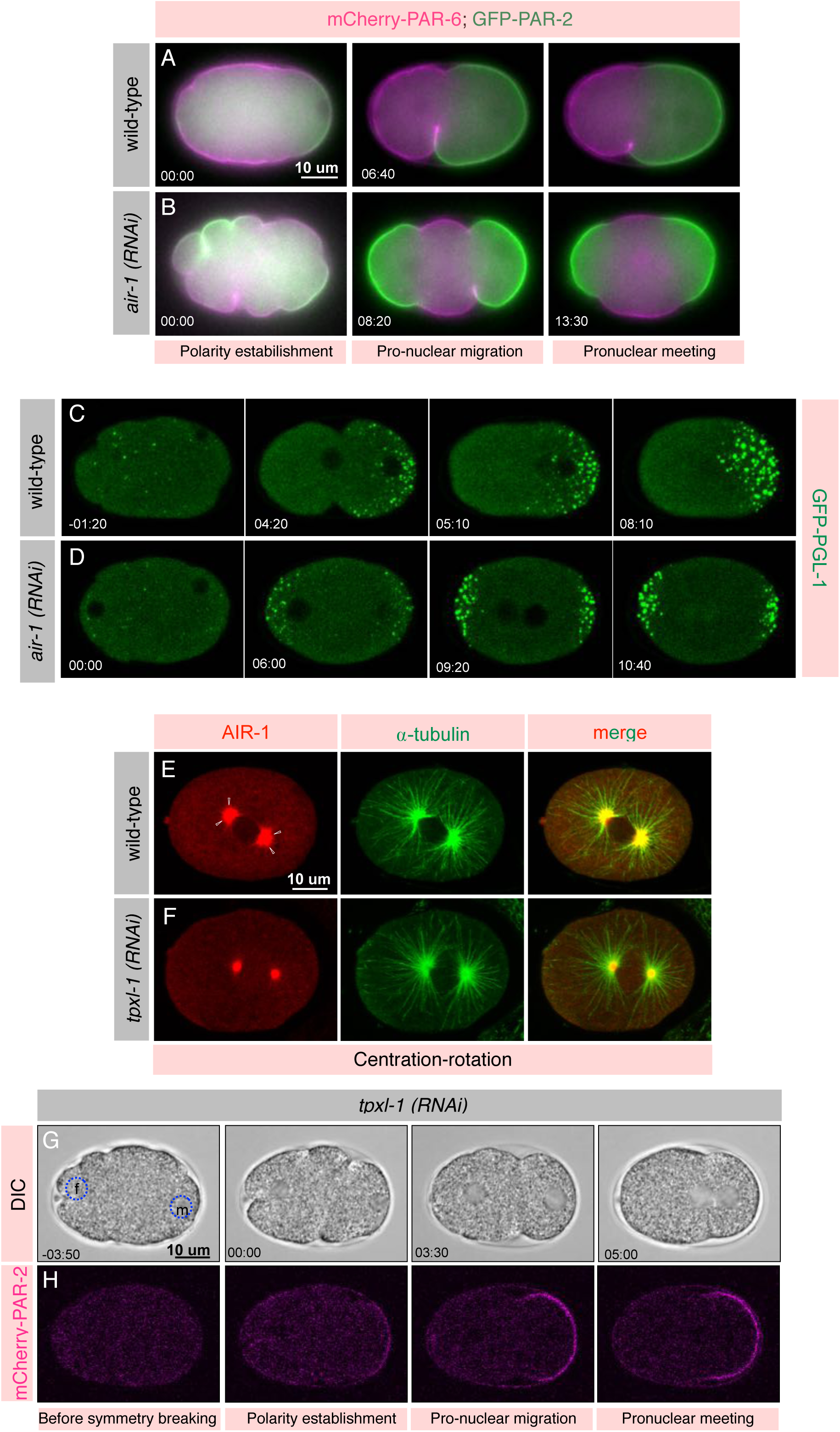
AIR-1 localization at the astral microtubules is not required for its impact on polarity establishment. (A, B) Images from time-lapse epifluorescence microscopy of embryos expressing mCherry-PAR-6 and GFP-PAR-2 in the wild-type (A) or in *air-1 (RNAi)* (B) at the various stages in the early embryos. See also corresponding Supplementary Movies S11 and S12. mCherry-PAR-6 signal is shown in magenta and GFP-PAR-2 is in green. Note the presence of excess membrane ruffles in the *air-1(RNAi)* embryo in comparison with the wild-type embryo. Also, note the appearance of two PAR-2 domains at the anterior and posterior cell cortex in the *air-1 (RNAi)* embryo in comparison with the single posterior PAR-2 domain in the wild-type embryos. More than 20 embryos were recorded and the represented embryos are shown. (C, D) Images from time-lapse epifluorescence microscopy of embryos expressing the P-granule component GFP-PGL-1 in wild-type (C) or in *air-1 (RNAi)* (D). GFP-PGL-1 signal is shown in green. See also corresponding Supplementary Movies S15 and S16. Note the presence of two GFP-PGL-1 enriched regions in *air-1 (RNAi)* embryos in comparison to the wild-type embryos. More than 10 embryos were recorded and the represented embryos are shown. (E, F) wild-type (E) or *tpxl-1 (RNAi)* (F) embryos either at the centration rotation stage stained for AIR-1 (red) and α-tubulin (green). Note the enrichment of AIR-1 at the astral microtubules (shown by white empty arrow head) that is absent in the *tpxl-1 (RNAi)* background. More than ten embryos were scored for each condition, and representative images are shown. (G, H) Images from time-lapse confocal microscopy in combination with DIC microscopy of embryos expressing mCherry-PAR-2 in wild-type (G) or in *tpxl-1 (RNAi)* (H) at the various stages in the early embryos. See also corresponding Supplementary Movie S17. mCherry-PAR-2 signal is shown in magenta. Note the collapsing of the mitotic spindle in *tpxl-1 (RNAi)* in the Supplementary Movie S17 as mentioned in the Ozlu et al., 2005. More than 10 embryos were recorded, and the represented embryos are shown.

**Supplementary Figure 3:**
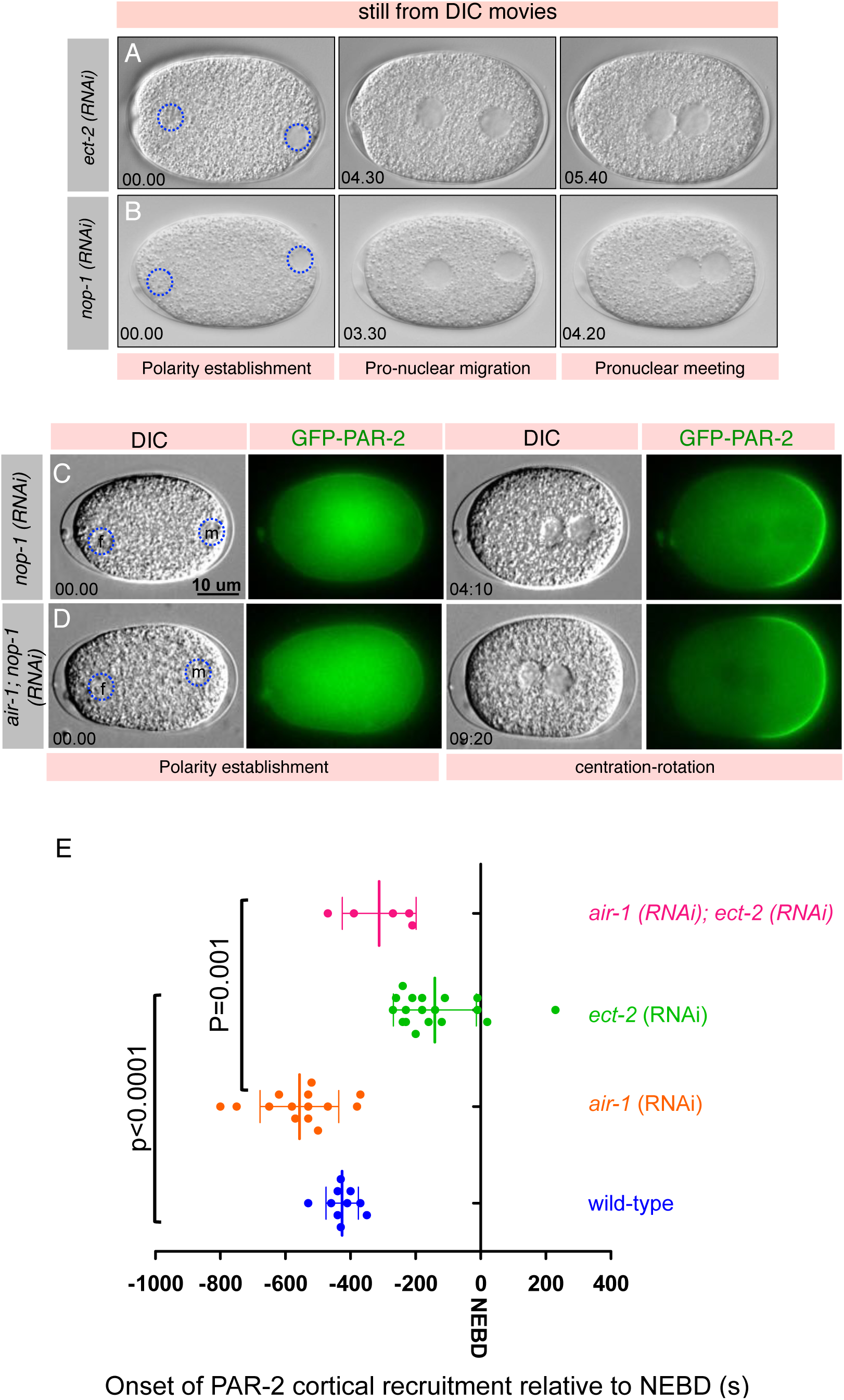
Loss of ECT-2 or its activator NOP-1 cause diminish cortical contractility in the one-cell embryo. (A, B) Images from time-lapse DIC microscopy of *C. elegans* early embryos in *ect-2 (RNAi)* (A) or *nop-1(RNAi)* (B). Time is shown in minutes which is solely based on the size of the male pronucleus because of the absence of cortical contractility in *ect-2 (RNAi)* or *nop-1 (RNAi)*. t=0 corresponding to the size equivalent of 5-6 μm. Ten embryos were analyzed in each condition and represented embryos are shown. Note loss of cortical contractility in *ect-2 (RNAi)* and *nop-1 (RNAi).* (C, D) Images from time-lapse epifluorescence microscopy in combination with DIC microscopy of embryos expressing GFP-PAR-2 in *nop-1 (RNAi)* (C) or *air-1 (RNAi); nop-1 (RNAi)* (D) at various stages as mentioned. GFP-PAR-2 signal is shown in green. Note that *nop-1 (RNAi)* and *air-1; nop-1 (RNAi)* shows the appearance of the posterior PAR-2 domain that is significantly later than wild-type as reported earlier (Tse et al., 2012). More than 10 embryos were recorded, and the represented embryo are shown. Quantification of the timing of the cortical localization of mCherry-PAR-2 in wild-type, *air-1 (RNAi), ect-2 (RNAi)* and *air-1 (RNAi); ect-2 (RNAi)*. Please note a significant delay in PAR-2 recruitment in *ect-2 (RNAi)* or *air-1 (RNAi); ect-2 (RNAi)* in contrast to the wild-type or *air-1 (RNAi)*. Also note that mitotic onset as monitored by nuclear envelope breakdown (NEBD) is delayed in *air-1 (RNAi)* as reported earlier in Hachet et al., 2007 and Portier et al., 2007. Time is reported in seconds (s).

